# Novel Conformation Specific Inhibitors of Activated GTPases reveal Ras-dependency of Patient-Derived Cancer Organoids

**DOI:** 10.1101/548750

**Authors:** Svenja Wiechmann, Pierre Maisonneuve, Britta M. Grebbin, Meike Hoffmeister, Manuel Kaulich, Hans Clevers, Krishnaraj Rajalingam, Igor Kurinov, Henner F. Farin, Frank Sicheri, Andreas Ernst

**Author notes:** Corresponding author: Andreas Ernst.

## Abstract

The small GTPases H, K, and NRAS are molecular switches that are indispensable for proper regulation of cellular proliferation and growth. Mutations in this family of proteins are associated with cancer and result in aberrant activation of signaling processes caused by a deregulated recruitment of downstream effector proteins. In this study, we engineered novel variants of the Ras-binding domain (RBD) of the kinase CRAF. These variants bound with high affinity to the effector binding site of active Ras. Structural characterization showed how the newly identified mutations cooperate to enhance affinity to the effector binding site compared to RBDwt. The engineered RBD variants closely mimic the interaction mode of naturally occurring Ras effectors and as dominant negative affinity reagent block their activation. Experiments with cancer cells showed that expression of these RBD variants inhibits Ras signaling leading to a reduced growth and inductions of apoptosis. Using the optimized RBD variants, we stratified patient-derived colorectal cancer organoids according to Ras dependency, which showed that the presence of Ras mutations was insufficient to predict sensitivity to Ras inhibition.

## Introduction

The small GTPases H, K, and NRAS are molecular switches that relay signals from growth factor receptor tyrosine kinases to transcription factors and other intracellular mediators to affect the growth, proliferation, and survival of cells. To achieve this function, the conformation of Ras GTPases cycles between an inactive, guanosine 5’-diphosphate (GDP)-bound state and an active, guanosine 5’-triphosphate (GTP)-bound state that interacts with downstream effector proteins. Along this reaction cycle, the weak hydrolysis activity of Ras is enhanced by GTPase activating proteins (GAPs), and guanine exchange factors (GEFs) facilitate the discharge of GDP and the reloading with GTP. In cancer, mutations in members of the Ras family shift the fine-tuned equilibrium of this reaction cycle toward the active, GTP-bound state, resulting in constitutive activation of downstream kinases. The resulting uncoupling of the regulatory link between proliferation and upstream receptor signaling leads to uncontrolled growth and proliferation. Thus, mutated Ras GTPases are oncogenic drivers in various malignancies making them, and their downstream effector kinases, a major focus for the development of anti-proliferative drugs.

When in active conformation, Ras GTPases propagate signals by the recruitment of kinases of the Raf family, thereby stimulating growth factor signaling pathways[1]. Members of this family of kinases interact with several different GTPases of the Ras subfamily. For example, BRAF is activated by oncogenic Ras and by the GTPase Rap1 [2, 3]. In a two-hybrid assay, the kinases ARAF and CRAF interact with the closely Ras-related GTPase RRAS [4]. Furthermore, the CRAF Ras binding domain (RBD) crystallized in complex with either HRAS or RAP1A, indicating that both interactions are sufficiently strong to yield co-crystal structures [5, 6]. This multi-specificity of Raf kinases relates to the conservation of the effector binding site and RBD at the atomic molecular level[7]. Thus, reagents that interfere with the Ras/Raf interaction are likely to efficiently inhibit the activation of growth signals over a wide range of conditions.

Developing inhibitors of Ras has been challenging because of the high affinity of guanosine di/tri-phosphate (GDP/GTP) for the GTP binding pocket of the Ras family of proteins and due to the flat topology of the conserved effector binding site [8, 9]. Consequently, to date, only a few small molecule inhibitors exist that directly interfere with Ras function [10]. To overcome the paucity of small molecules that directly interfere with Ras activity, several affinity reagents based on protein scaffolds have been developed to address different facets of Ras biology. For example, fibronectin-derived monobodies inhibit signaling by disrupting the dimerization of H and KRAS at the plasma membrane [11]. Interfering with Ras dimerization limits the growth of xenografts that inducibly express the monobody [12]. An alternative strategy for inhibiting Ras signaling with engineered proteins is to prevent the recruitment of downstream effectors or block the activation of Ras by GEFs. To this end, affinity reagents have been generated that compete with Ras effectors by selectively binding to the active or inactive state of oncogenic Ras [13-19]. However, several required high concentrations to be effective in cells [16, 17, 20], and their selectivity for members of the Ras family is unknown. The limited effectiveness may be related to the interaction of Raf family kinases with various Ras GPTases. Thus, a rationale for affinity reagents with improved efficacy could be those that block interactions with multiple Ras family members simultaneously.

An ideal affinity reagent that inhibits Ras function should avoid steric constraints by precisely replicating effector binding to activated, GTP-bound Ras. Additionally, these affinity reagents should have a high affinity to the effector binding site and the ability to outcompete downstream effectors. Consequently, such an affinity reagent would act as inhibitor of Ras signaling, which could be used to probe the biology of Ras inhibition in cellular and patient-derived model systems. To derive an affinity reagent that fulfills these criteria, we adapted our strategy of optimizing pre-existing intermolecular contacts [21, 22] to the RBD of CRAF. We constructed a phage-displayed RBD library encoding mutations of interface residues and selected for improved binding to active, GTP-bound Ras. Crystal structures of complexes with activated HRAS G12V showed that the RBD variants (RBDvs) precisely mimic effector binding, and the engineered mutations subtly improve intermolecular contacts. Importantly, intracellular expression of individual RBDvs resulted in impaired growth of various cancer cell lines due to a robust inhibition of Ras signaling and the MAP kinase signaling cascade. Finally, in patient-derived organoids, the differential effects of the RBDvs on cellular growth and metabolic activity revealed differences in Ras dependency of colorectal tumor organoids, demonstrating that genetic data were insufficient to predict responsiveness to Ras inhibition. The ability of the RBDvs to inhibit growth of cells lacking activating mutations in Ras family members may relate to their multispecificity in interacting with active conformations of Ras GTPases with a conserved switch 1 region.

## Results

### Engineered RBD variants have high affinity for GTP-bound Ras and compete with effector binding *in vitro*

Previous studies demonstrated that the CRAF-RBD is highly tolerant to mutations and can be computationally engineered to preferentially bind inactive states of Ras [18, 23, 24]. To optimize the interface of the RBD to active Ras, we analyzed existing crystal structures of Ras:RBD complexes [5, 25, 26]. In total, we identified 14 residues that make sidechain contacts to the switch-1 region of the Ras effector binding site. These residues are located on the β1-β2 hairpin and α1 helix, and span two distinct regions of the CRAF-RBD. To achieve a moderate to low mutation rate that does not alter the binding mode nor impair the structure of the RBD, we used a soft randomization strategy that allowed 70% of the wild-type nucleotides and 10% of the non-wild-type nucleotides to occur at any codon position encoding the targeted residues (**Fig. 1A**). Additionally, we replaced three unpaired cysteines at position 81, 95, and 96 of the RBD with serine residues to prevent dimerization of phage-displayed proteins and improve overall presentation of the RBDvs. Our final library contained more than 2×10^9^ unique RBDvs presented on the surface of filamentous phage and subsequent selection by phage display yielded 5 variants with improved binding to surface immobilized GTPγS-loaded HRAS (**Fig. 1B**). The selected variants show a conserved mutational pattern, indicating a shared binding mode to activated Ras. Strikingly, the mutation Gln^66^ to Glu is highly enriched in all 5 variants, indicating that this mutation has a crucial role in mediating the increased affinity. Coincidentally, the mutation from Val^88^ to Arg, previously shown to increase the affinity of RBD to HRAS [27], was also present in all 5 variants. An Arg^89^ to His mutation occurred in four of the five RBDvs.

**Figure 1.**
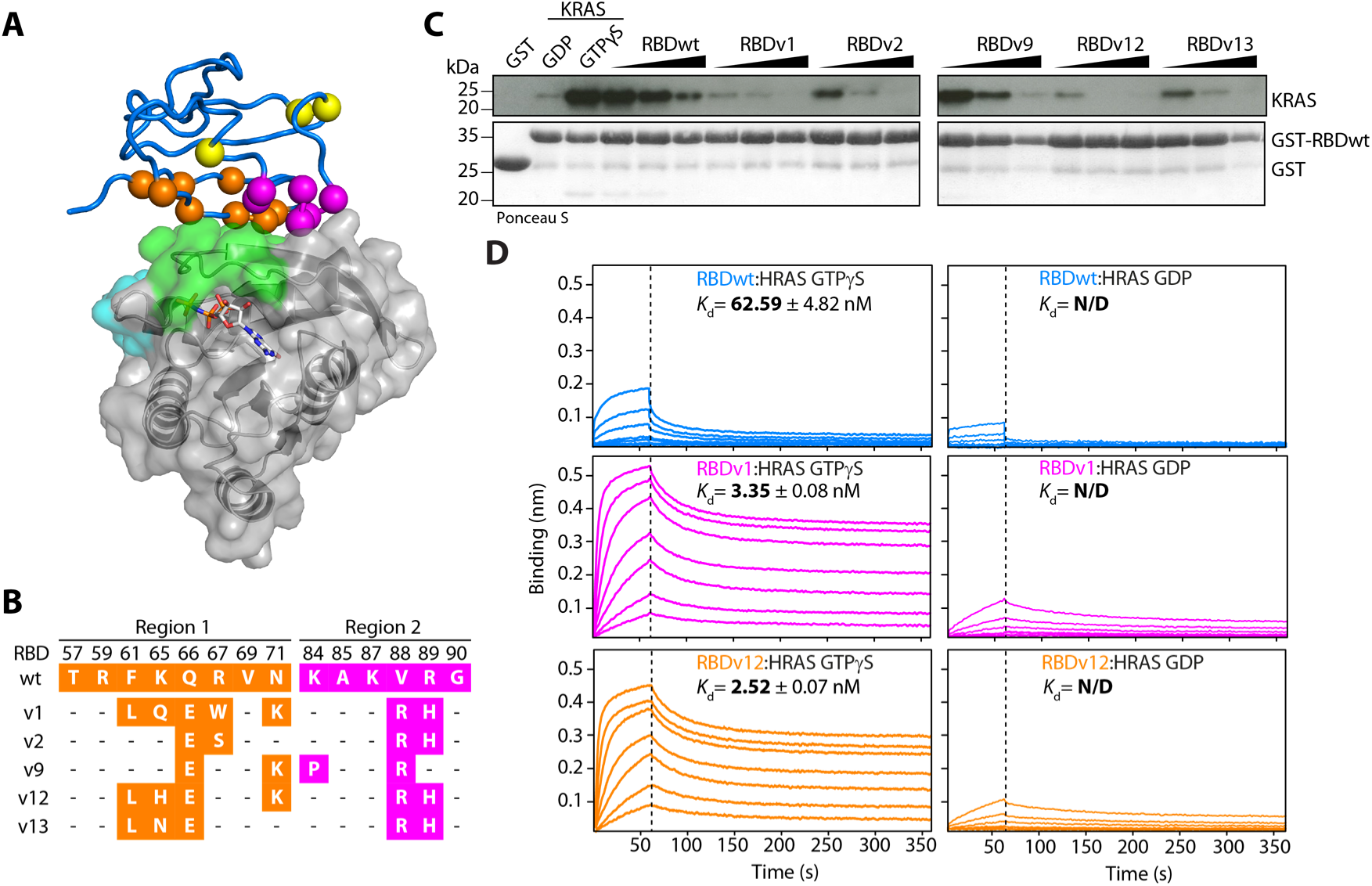
CRAF-Ras-binding domain (RBD) library design and selected RBD variants (RBDvs) outcompete RBDwt by binding with highly improved affinity to active Ras. (**A**) CRAF-RBD in complex with HRAS (pdb: 4G0N) [5]. The RBD is shown as blue tube and HRAS as transparent surface representation indicating the location of switch 1 (green) and switch 2 (cyan). Residues modified in region 1 (orange), region 2 (magenta) and Cys to Ser mutations (yellow) of CRAF-RBD are shown as colored spheres. (**B**) Sequences of RBDvs selected by phage display. Residues in region 1 and region 2 are colored as in (A). Non-mutated positions are indicated by dashes. (**C**) *In vitro* competition of His-tagged GTPγS-loaded KRAS binding to GST-tagged RBDwt immobilized on glutathione sepharose resin with increasing molar ratios of His-tagged RBDvs or RBDwt (1:1, 1:2.5 and 1:10). KRAS bound to beads was detected by immunoblot and the corresponding Ponceau S stained membrane is shown. (**D**) Binding of GTPγS- and GDP-loaded HRAS to immobilized GST-tagged RBDwt (blue), RBDv1 (magenta) or RBDv12 (orange) measured by bio-layer interferometry (BLI). Concentrations of Ras ranged from 1 µM to 15.6 nM in a 1:1 dilution series. *K*_d_ values for each experiment are shown.

After purification as His-tagged proteins, we tested if the engineered RBDvs outcompeted CRAF-RBDwt binding to GTPγS-loaded KRAS *in vitro* (**Fig. 1C**). Relative to RBDwt, all engineered variants showed an enhanced ability to compete off GTPγS KRAS in solution. A similarly enhanced competitive ability of the RBDvs was observed using the Ras association domain (RA) of Ral Guanine Nucleotide Dissociation Stimulator (RalGDS) instead of the CRAF-RBDwt (**Supplementary Fig. 1A**). Because RBDv1 and RBDv12 performed best in these experiments, we focused our further analysis on these two variants.

Affinity measurements confirmed that RBDv1 and RBDv12 have improved binding properties relative to RBDwt for activated HRAS with K_d_s of ∼ 3 nM versus ∼60 nM and strongly preferred binding to activated, GTPγS loaded HRAS with negligible binding to the GDP-bound form (**Fig. 1D**). In control experiments, we confirmed that His-tagged HRAS GTPγS does not bind to GST-loaded sensors or empty sensors alone (**Supplementary Fig. 1B**), indicating that the slow off rate is not due to non-specific binding of HRAS to GST or the sensor material.

### The engineered RBDvs mimic effector binding to active Ras

To understand the structural basis for improved binding, we crystallized HRAS G12V in complex with RBDv1 at 2.9 Å and RBDv12 at 3.15 Å resolution in an active conformation (**Fig. 2A, Supplementary Table 1**). Structural analysis revealed that the RBDvs engaged the effector binding site of HRAS through a canonical binding mode [5] with only minor shifts to the center of mass positions and rotation angle (1.8 Å/15° and 2.0 Å/16° for RBDv1 and RBDv12, respectively). Similar to the canonical binding mode of RBDwt, β2 of the RBDvs forms an extended intermolecular β-sheet with β2 of HRAS (**Supplementary Fig. 2A**). In addition, helices α1 of both RBDwt and RBDvs form direct contacts with switch 1 residues in Ras (**Fig. 2A**). Inspection of direct contact interface reveals that the conserved RBDvs mutations of Gln^66^ to Glu, Val^88^ to Arg, and Arg^89^ to His result in a rewiring of the hydrogen (H)-bonding pattern to the Asp^38^ and Tyr^40^ side chains in HRAS (**Fig. 2B**). This change in H-bonding pattern together with steric effects involving Ile^21^ in HRAS and Val^88^ to Arg in RBDvs appears responsible for a shift of the α1-helix of the RBDvs relative to that observed in RBDwt (**Fig. 2C**). In summary, our combinatorial approach to RBD-interface engineering identified three key mutations that work together to improve contacts to residues in the switch 1 region of Ras in an active conformation. Because the binding mode of RBDvs to HRAS overlaps completely with RBDwt, the RBDvs are expected to outcompete binding of Ras effectors that engage this common binding site.

**Figure 2.**
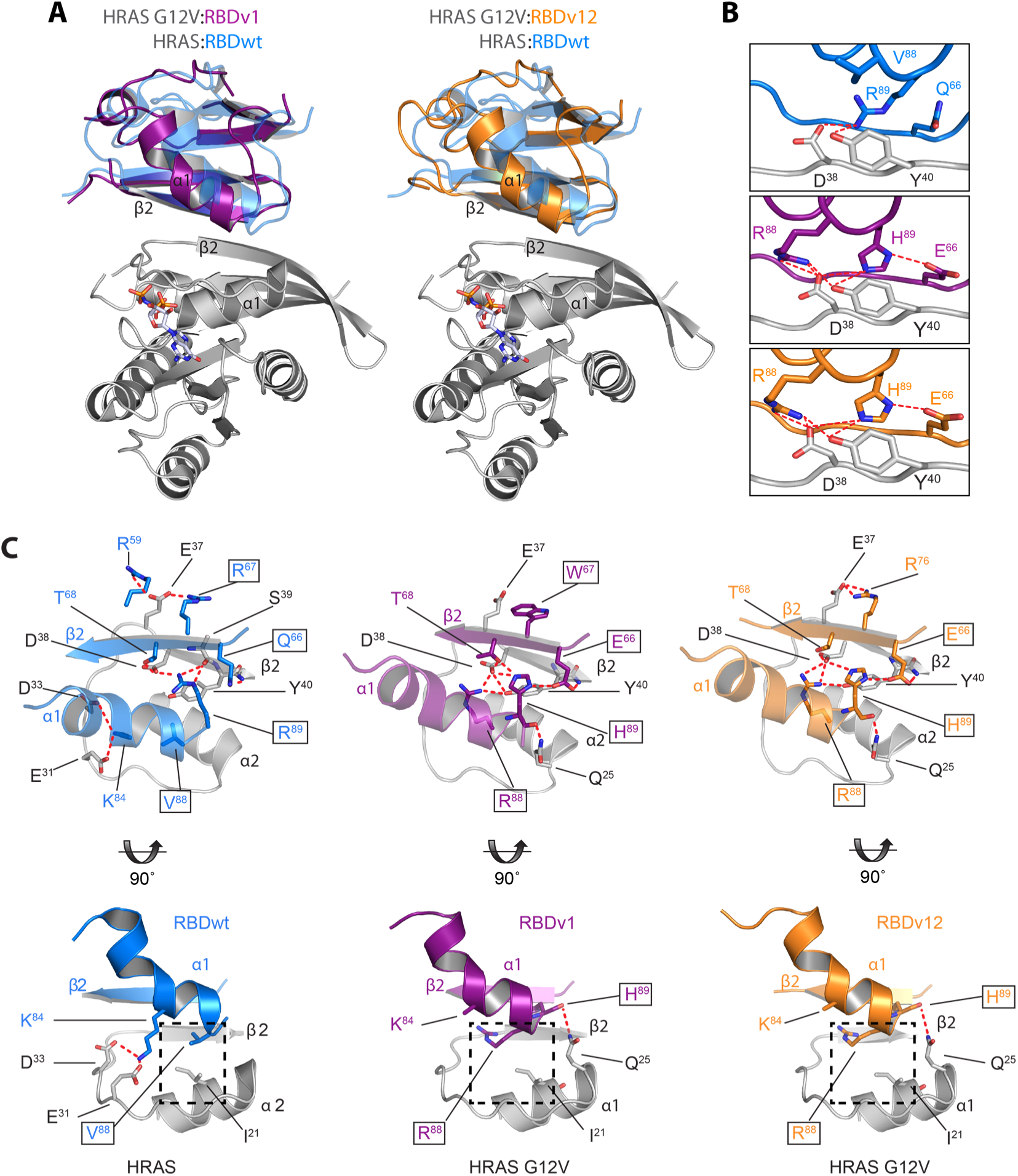
RBDvs bind at the Ras effector binding site. (**A**) Crystal structures of HRAS G12V (grey) in complex with RBDv1 (magenta) at 2.9 Å or RBDv12 (orange) at 3.15 Å resolution. Both structures are superimposed on HRAS in complex with RBDwt (blue) (pdb: 4G0N) [5]. For clarity, only HRAS G12V and the GTP analog phosphoaminophosphonic acid-guanylate ester (GNP) is shown as colored sticks. (**B**) Side by side comparison of HRAS in complex with RBDwt (upper panel), RBDv1 (middle panel) or RBDv12 (lower panel) showing changes caused by mutations at positions 66, 88 and 89. Polar interactions are indicated by dashed lines (red). HRAS G12V, RBDwt, RBDv1 and v12 main and side chains are colored as in (A). Residues are numbered according to the PDB entry for 4G0N. (**C**) Top and side view of the binding interface of RBDwt and RBDvs with HRAS. Residues involved in intermolecular interactions are shown as stick. Residues that are mutated in RBDvs are highlighted by a black square. 90° rotated side view highlights the steric clash between Ile^21^ in HRAS and Val^88^ to Arg in RBDvs that is involved in a shift of the α1-helix of the RBDvs relative to that observed in RBDwt.

### The RBDvs bind Ras GTPases in cells

We tested the intracellular specificity of the RBD variants using mass spectrometry. We immunoprecipitated hemagglutinin (HA)-tagged RBDvs and HA-RBDwt that were inducibly expressed in stably transduced colon carcinoma HCT 116 cells (**Supplementary Fig. 3A and B**). HCT 116 are heterozygous for the activating mutant KRAS G13D. Strikingly, relative to RBDwt, RBDv1 and RBDv12 displayed up to 5500-fold enrichment of peptides from endogenous KRAS4B G13D isoform (**Fig. 3A, Supplementary Table 2**). The prevalence of peptides originating from the constitutively active KRAS4B G13D isoform suggests that the RBDvs preferentially interact with Ras GTPases, which are in an active conformation.

**Figure 3.**
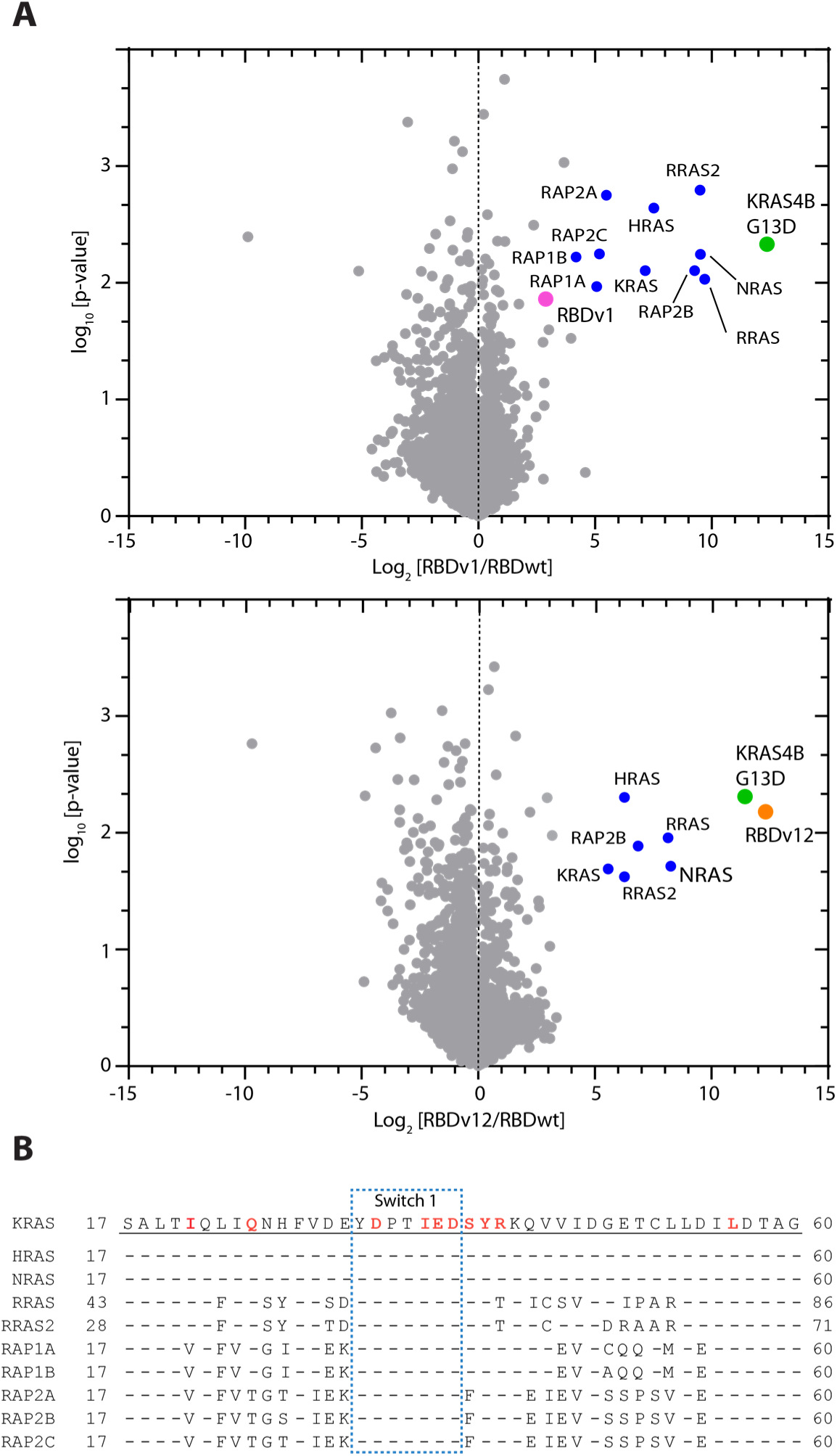
RBDvs are specifically binding to endogenous KRAS4B G13D in HCT 116 cells. (**A**) Mass spectrometry of co-immunoprecipitation using anti-HA beads from lentiviral transduced HCT 116 cells stably expressing HA-tagged RBDwt, RBDv1 and RBDv12 upon induction with doxycycline (DOX) (1 µg/ml, 24h). Detected proteins (grey) were plotted as log_2_ average protein intensity for RBDv1/RBDwt (upper panel) or RBDv12/RBDwt (lower panel) versus _log10_ p-value. Only proteins that have been identified with a p-value < 0.05 are shown (n=2). More than 16-fold enriched proteins and the RBDvs are shown as indicated (colored symbols). (**B**) Sequence alignment of Ras GTPases that have been enriched more than 16-fold (log2 > 4). Conserved residues relative to KRAS are indicated as dashes. Switch 1 residues (blue box) and residues within 4.5 Å of RBDvs (magenta) are highlighted.

We also detected peptides from other Ras family GTPases, although these were ∼10-fold less abundant than KRAS4B G13D peptides. Analysis of the primary sequences of the other Ras family members that interacted with the RBDvs showed that these Ras family members share a common effector binding site (**Fig. 3B**). In particular, the detected peptides were from Ras family members containing D^38^ and Y^40^ residues in their respective switch-1 region, which we identified as key interaction residues for the RBDvs (**Fig. 2B**). Thus, the immunoprecipitation data showed that RBDvs bind to their intended targets, Ras proteins, in cells, and exhibit the highest affinity for active Ras.

Consequently, these results suggested that the RBDvs are dominant-negative affinity reagents that will not only impair signaling through constitutive active Ras mutants but also other Ras GTPases in the active conformation. Thus, the RBDvs have the potential to inhibit multiple Ras-mediated pathways to Raf activation. Furthermore, the simultaneous inhibition of multiple Ras GTPases is potentially an efficient way to control the proliferation of cells with heterozygous KRAS genotypes, which confer resistance to MEK inhibitors [28].

### RBDvs inhibit activation of the ERK and AKT pathways resulting in reduced metabolic activity and apoptosis

To test if the RBDvs act as inhibitors of Ras-dependent signaling processes, we probed whole cell lysates from cell lines expressing HA-tagged RBDvs or RBDwt with antibodies detecting the phosphorylation state of kinases downstream of Ras proteins. Specifically, we examined the effect of inhibition of Ras/Raf interaction in cell lines of different cancer origins and with different Ras mutations (**Supplementary Table 3**). Immunoblot analysis showed that the RBDvs reduced the phosphorylation of the growth signal-activated ERK1/2 in HCT 116, Mia PaCa-2, A549, and H1299 cells (**Fig. 4A**). The corresponding control experiments with RBDwt, despite its higher expression, did not reduce ERK1/2 phosphorylation, indicating that the improved affinity of the RBDvs for active Ras isoforms is required to suppress activation of these downstream kinases.

**Figure 4.**
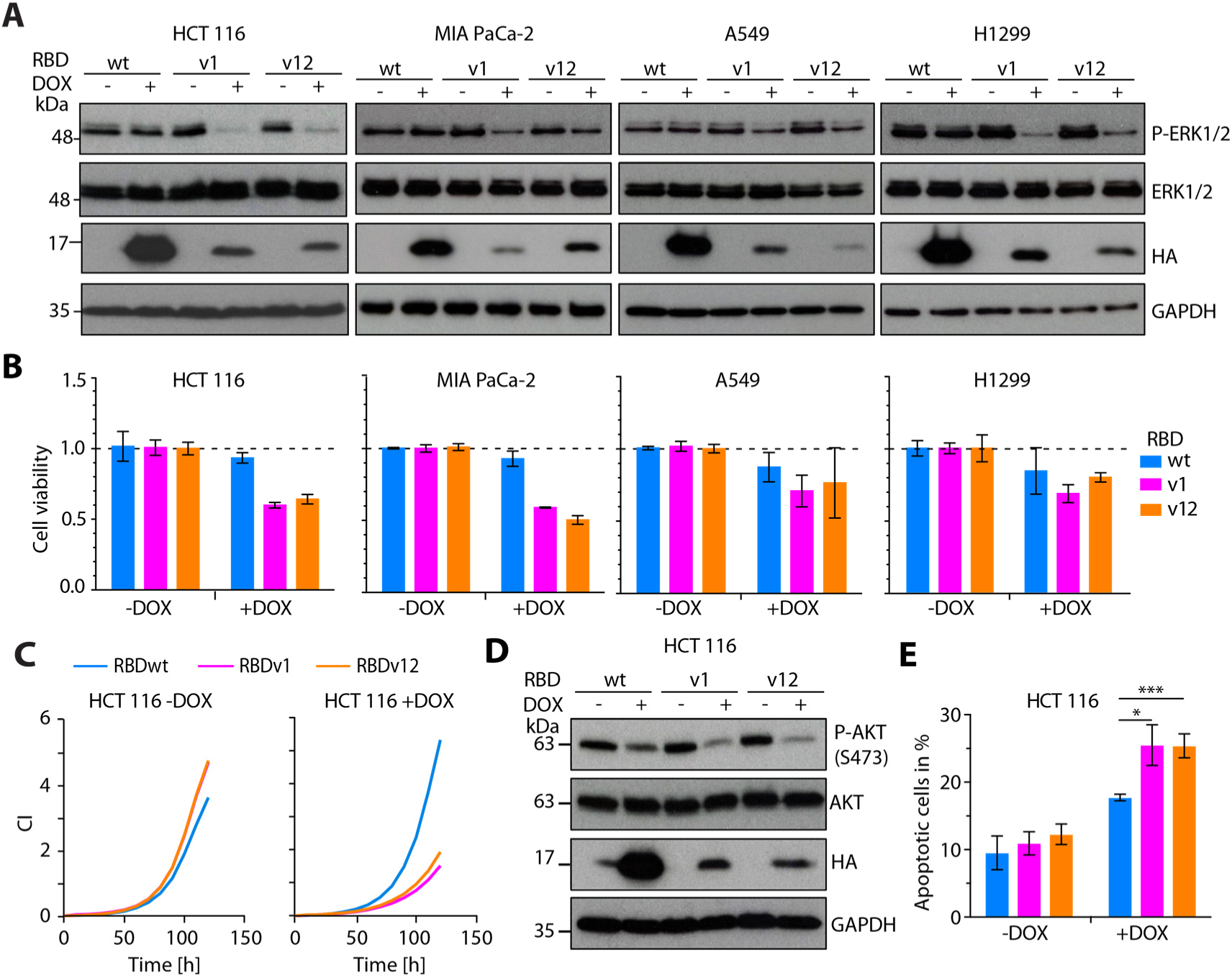
RBDvs inhibit MAPK and PI3K signaling in cancer cells, reduce cellular viability and induce apoptosis in HCT 116 cells. (**A**) Immunoblot of whole cell lysates from HCT 116, MIA PaCa-2, A549 and H1299 stably transduced with inducible lentiviral constructs expressing HA-tagged RBDwt, RBDv1 or RBDv12 in absence (−) and presence (+) of DOX (1 µg/ml, 24 h). Cell lysates were analyzed using the indicated antibodies. (**B**) Normalized cell viability from cells used in panel (A) transduced with RBDwt (blue), RBDv1 (magenta) or RBDv12 (orange) measured as cellular ATP content by luciferase mediated bioluminescence. Reduction of cellular ATP in presence of RBDs (+DOX) was monitored after 120 h induction and normalized to the luminescence of non-induced control (-DOX) cells. (Error bars corresponds to ± standard deviation (SD) of 3 biological replicates (n=3). (**C**) Cellular growth measured following cell index (CI) over time (h) by Real-time cell analysis (RTCA) of HCT 116 cells in absence or presence of DOX (1 µg/ml, 120 h). The mean of two technical replicates is shown. (**D**) Immunoblot of whole cell lysates from HCT 116 stably transduced with RBDwt, RBDv1 or RBDv12 in absence (−) and presence (+) of DOX (1 µg/ml, 24 h). Cell lysates were analyzed using the indicated antibodies. (**E**) Quantification of flow cytometry data of annexin V antibody and propidium iodide (PI) stained HCT 116 cells cultured under full medium conditions in absence (-DOX) or presence (+DOX) (1 µg/ml, 72 h). (Error bars corresponds to ± standard deviation (SD) of 3 biological replicates (n=3). P-values were calculated by an unpaired t test (\**P*< 0.05, *P< 0.01, \*\*\**P*< 0.005).

Reduced activity of MAP kinase pathway often results in reduced cell viability. Therefore, we measured ATP content, as an indicator of metabolic activity and viability, and monitored growth curves, as an indicator of proliferation. In all 4 cell lines, inhibition of effector binding to active Ras by the RBDvs reduced cell viability (ATP content) (**Fig. 4B**), indicating that the RBDvs disrupt Ras signaling in cells from different cancer backgrounds. Importantly, RBDwt control had only minimal effects on the viability of all 4 cell lines. Five-day growth curves measured with HCT116 cells confirmed that the RBDvs, but not RBDwt, inhibited proliferation (**Fig. 4C**).

Ras signaling not only mediates proliferative responses, but these GTPases also promote cell survival. In the presence of the RBDvs, HCT 116 cells exhibited reduced phosphorylation of Ser^473^ in the serine/threonine kinase AKT, indicating that PI3 kinase pathway is inhibited (**Fig. 4D**). Together, the reduced MAP kinase and PI3 kinase pathway activity could not only reduce cellular metabolic activity and proliferation but could also increase apoptosis. We monitored HCT116 cells expressing either RDBvs or RBDwt for annexin V staining as an indicator of apoptotic cells (**Fig. 4E, Supplementary Fig. 4**). Quantification of annexin V staining by flow cytometry revealed that HCT116 cells expressing the RBDvs had significantly increased number of apoptotic cells compared to non-induced controls and compared to induced cells expressing RBDwt. In summary, these results showed that the RBDvs inhibit the ERK and PI3K signaling pathway, resulting in growth reduction in a wide range of cancer cell lines and inducing apoptosis in HCT116 cells.

### RBDvs lead to reduced growth in patient-derived colorectal cancer organoids

To investigate whether the characteristics of our RBDvs in cell culture can be translated into a patient-derived model, we used tumor organoids with known Ras mutation status isolated from surgically removed colorectal carcinoma from 7 patients [29] (**Table 1**). After transduction with the doxycycline-inducible lentiviral constructs, we compared cell viability and growth of organoids, cultured in Matrigel and expressing RBDvs or RBDwt. We evaluated by immunoblot of organoid lysates ERK and AKT phosphorylation in response to doxycycline-induced expression of the RBDvs or RBDwt (**Fig. 5A, Supplementary Figure 5A**). The different organoids showed different sensitivity to the inhibitory effects of the RBDvs on these two kinases. For example, P17T with wild-type KRAS showed little effect of RBDv12, and P6T with KRAS G12C showed a stronger effect of the RBDvs on ERK1/2 phosphorylation than on Akt phosphorylation (**Fig. 5A**). P18T with wild-type KRAS showed little effect of either RBDv1 or RBDv12 (**Supplementary Figure 5A**). This is consistent with different cancers having diverse adaptive signaling pathways and different signaling dependencies.

**Table 1:** Summary of response to Ras inhibition and mutational status of tested CRC organoids P: patient; T: tumor; MSI: Microsatellite instability.

| HUB tumor organoid[29] | Organoid response in presence of RBDvs |  |  |  | Published mutational status[29] |  |  |  |  |
| --- | --- | --- | --- | --- | --- | --- | --- | --- | --- |
|  | pERK | pAKT | Cell Viability | Colony growth | Geftinib sensitivity | KRAS | NRAS | BRAF | PI3K |
| P6T | ↓ | – | ↓ | ↓ | resistant | G12C | wt | wt | wt |
| P26T | – | – | – | – | resistant | G12V | wt | wt | wt |
| P28T | ↓ | ↓ | ↓ | ↓ | sensitive | G12V | wt | wt | wt |
| P17T | ↓ | ↓ | ↓ | ↓ | sensitive | wt | wt | wt | wt |
| P18T | – | – | – | – | resistant | wt | wt | wt | wt |
| P20T | – | – | – | – | sensitive | wt | wt | wt | wt |
| P24bT (MSI) | ↓ | ↓ | ↓ | ↓ | n/a | wt | wt | wt | wt |

**Figure 5.**
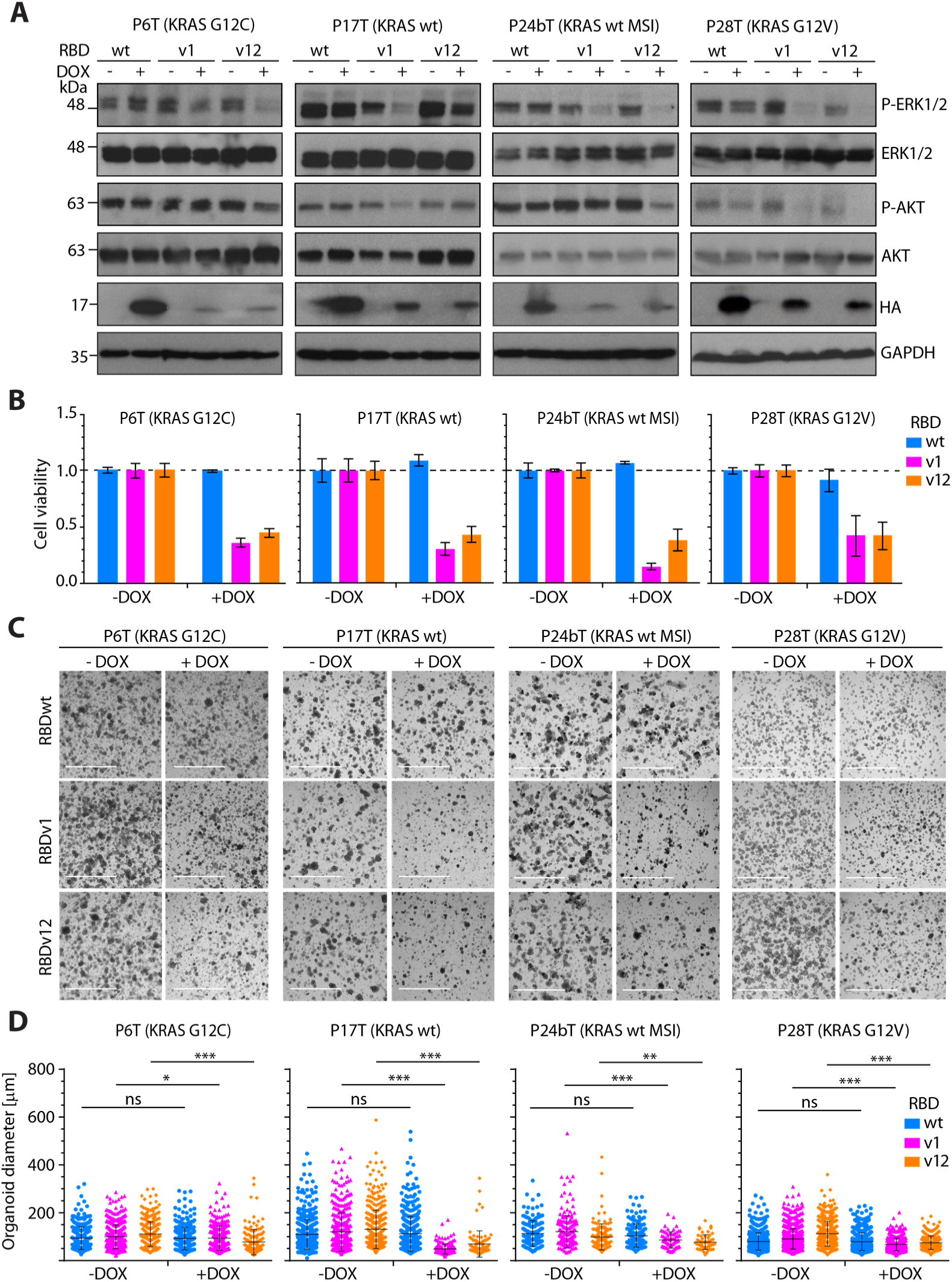
Patient-derived colorectal cancer (CRC) organoids show reduced MAPK and AKT signaling and reduced cell viability upon RBDvs expression. (**A**) Immunoblot of whole cell lysates derived from indicated patient-derived CRC organoids stably transduced with lentivirus encoding HA-tagged RBDwt, RBDv1 and RBDv12 in absence (−) or presence (+) of DOX (2 µg/ml, 72 h). Cell lysates were analyzed using indicated antibodies. (**B**) Cellular ATP content of organoid cultures used in (A) expressing RBDwt (blue), RBDv1 (magenta) and RBDv12 (orange) was measured in a luciferase mediated bioluminescence assay. Reduction of cellular ATP in presence of RBDvs (+DOX) in P17T and P24bT was monitored after 72 h induction and normalized to the luminescence of non-induced control organoids (-DOX) (2 µg/ml, 72 h). P6T and P28T stable organoids were seeded as single cells and induced for 96 h and analyzed as above. Error bars correspond to ± SD of three technical replicates (n=3). (**C**) Bright field microscope images of organoid cultures used in (A) in presence (+) or absence (−) of DOX (2 µg/ml, 72 h). Scale bars correspond to 2 mm. P=patient; T=tumor, MSI=microsatellite instability. (**D**) Quantification of organoid size in presence and absence of DOX (2 µg/ml, 2-6 d) from bright field microscope images of organoid lines used in (A). Error bars correspond to ± SD. Statistics were calculated using the Mann-Whitney U-test (\**P*< 0.05, \*\**P*< 0.01 and \*\*\**P*< 0.005).

Despite the variability in the effects on kinase phosphorylation, subsets of the organoids with mutant Ras (P6T and P28T) and those with wild-type Ras (P17T and P24bT) had reduced viability when expressing either RBDv1 or RBDv12, as indicated by reduced metabolic activity (ATP content) (**Fig. 5B**). In addition, the effect of RBDvs on cell viability correlated with decreased colony size as determined by bright field microscopy (**Fig. 5C**). Importantly, quantification of the colony size indicated that the inhibitory RBDvs significantly reduced growth when compared to non-induced organoids (**Fig. 5D**). However, growth of P18T and P20T, both with wild-type KRAS, and P26T with KRAS G12V was unaffected by either RBDv1 or RBDv12 (**Supplementary Figure 5B**). Thus, KRAS mutant status was insufficient to predict sensitivity to the growth-inhibiting effects of Ras inhibition (**Table 1**). Our data indicated that the RBDvs classify colorectal cancer samples for Ras dependency. The difference in responsiveness to RBDvs was not related to downstream activation of growth signaling by mutant BRAF or PI3K in these organoids, because we tested organoids that are wild-type for BRAF and PI3K[29]. In conclusion, the RBDvs decreased the growth of four patient-derived organoid lines, indicating that inhibiting the interactions between activated Ras proteins and their effectors may be a valid strategy in cancer therapy.

## Discussion

By engineering the Ras/Raf interface of the CRAF-RBD, we developed potent and highly selective inhibitors of activated Ras that outcompete the binding of signaling effectors. The selectivity of the engineered variants for an active conformation of Ras occurred through molecular contacts to a minimal epitope composed of switch 1 residues at the center of the effector binding site. High affinity was achieved by a subtle rewiring and optimization of the hydrogen bond pattern at the interface between the RBD and HRAS. Because the effector binding site is conserved amongst Ras family members, we detected interactions between the RBDvs and other Ras GTPases with nearly identical sequence composition in the switch 1 region. This multispecificity may be beneficial for therapeutic applications based on the RBDvs, because cancer cells often develop resistance to highly specific targeted therapies [30, 31]. The ability to inhibit multiple related Ras family members specifically in the active conformation may prevent a bypass or network rewiring that enables resistance, may interfere with other pathways that collectively provide cancer cells a growth advantage, or may sensitize cancer cells to other therapeutics.

In cellular experiments, expression of RBDvs efficiently reduced ERK and AKT phosphorylation and cellular growth and triggered apoptosis in cell lines from different cancer backgrounds. We applied the RBDvs to a clinical research question and showed that the RBDvs could be used as a tool to delineate Ras dependency in colorectal cancer carcinoma (CRC) [29]. In CRC several different signaling pathways have been implicated in disease pathogenesis [32]. However, the extent and heterogeneity of genetic alterations in CRC makes it difficult to analyze the contribution of individual pathways to the proliferative phenotype [29]. More importantly, this diversity makes defining an effective treatment strategy challenging. We showed that the inhibition observed in adherent cell culture experiments translates to patient-derived colorectal cancer organoids: Several of the organoids exhibited reduced growth when expressing either RBDv1 or RBDv12. However, indicative of their varying degree of Ras dependency, not all organoids responded equally well to Ras inhibition by the inhibitory RBDvs. Unexpectedly, the Ras dependency did not correspond to the presence of mutant Ras, showing that genetic information is insufficient to predict therapeutic response. Thus, the RBDvs can be used to facilitate functional classification of Ras dependency in intestinal tumor organoids, which has the potential to drive therapeutic strategies.

Although the RBDvs are a unique tool for studying Ras-dependent signaling processes, several intracellular affinity reagents targeting GTP-bound Ras have been reported [13, 15-17]. Compared with the previously reported Ras-targeted reagents, the RBDvs were effective at lower concentrations. For example, the affinity reagent R11.1.6, based on a DNA-binding domain from a thermophilic archaeon, binds Ras in an active conformation and competes with effector binding [15]. Although initial experiments in HEK293T cells suggested otherwise, R11.1.6 does not affect Ras-mediated signaling in a broad range of cancer cell lines [20]. Kauke *et al*. concluded that a higher concentration of R11.1.6 than was achieved by lentiviral transduction is required to efficiently outcompete Ras-binding effectors. Similar observations have been made for intracellular antibodies targeting the Ras-effector binding interface. The antibody fragment iDab#6 required the addition of a membrane localization peptide to overcome the binding avidity of endogenous Ras effectors to inhibit Ras-dependent signaling events [16]. The cell-penetrating TMab4 RT11 antibody targeting the switch 1 site of Ras proteins also requires high concentrations to effectively inhibit Ras-mediated signaling events [17]. Two designed ankyrin repeat proteins (DARPins) with specificity for either the GDP-bound inactive state (K27) or the GTP-bound active state (K55) of Ras have been reported [13]. Both DARPins inhibit Ras signaling in transfected HEK293T cells and lentivirus transduced HCT 116 cells; however, it remains to be seen if the observed effects occur in other cell types and organoids. The lower affinity of K55 (167 nM) compared with that of CRAF-RBDwt (∼ 60-80 nM) for GTP-bound Ras also suggests that a high concentration will be required to compete for endogenous effectors in a cellular context.

In contrast to these other Ras interaction inhibitors, the intracellular inhibition of Ras signaling by the RBDvs does not show similar avidity or concentration dependent effects. Indeed, we observed robust inhibition in several different cell lines and organoids despite the expression of the RBDvs was always less than the corresponding wild-type control. The RBDvs exhibited preferential binding to active Ras proteins, indicating high selectivity not only for Ras proteins with a shared effector binding site but also for these proteins in their active conformation. Most likely, the other interaction inhibitors, especially TMab4 RT11 that also binds the switch-1 sequence, may also have similar multispecificity for Ras proteins. This potential property of the other affinity reagents should be further examined.

A common challenge in all these efforts is the effective delivery of intracellular affinity reagents to the cytosol. Consequently, various intracellular protein delivery platforms, such as cell-penetrating peptides, nanocarrier, liposomes, polymer and nanoparticle-stabilized nanocapsules (reviewed in [33]), are actively under investigation. Another promising strategy for the delivery of proteins into mammalian cells is the use of bacterial toxins, which can deliver a wide array of passenger proteins spanning a range of sizes, structures, and stabilities[34]. For example, a recent report shows that a fusion protein of the RBDwt is able to pass through the channel of a tripartite toxin complex (Tc toxins) derived from bacterial pathogens. Thus, due to its small size and favorable charge distribution the RBD-scaffold may be particularly suited for delivery by Tc toxins [35]. In future, these and other platforms can enable targeted delivery of proteins into cells to realize the potential of protein-based therapeutics with intracellular sites of action. RBDvs would then be a candidate for delivery to test in treating diseases associated with unchecked Ras activity.

## Material and Methods

### Ras expression, purification and nucleotide exchange

The human HRAS (AA 1-166) and KRAS (AA 1-169, isoform B) proteins were expressed as GST-fusions from pGEX-6P-1 plasmids for selection experiments and as His-tag fusions from pET-53 plasmids for *in vitro* competition and binding assays. Plasmids were used to transform *E. coli* Rosetta(DE3) cells and individual colonies were handled essentially as written before [22], except that all buffers were supplemented with 5 mM MgCl_2_. Briefly, resulting cultures from individual colonies were inoculated into Luria Broth (LB) media and protein expression was induced at OD_600_=0.8 with 0.5 mM IPTG. After overnight incubation at 16 °C, bacterial pellets were resuspended in 50 mM Tris-HCl pH 7.5, 150 mM NaCl, 1 mM PMSF and 5 mM MgCl_2_ and lysed by sonication. The lysates were clarified by centrifugation and proteins in the supernatant were purified using Ni-NTA chromatography (Qiagen) for His-tagged proteins or Glutathione Sepharose 4 Fast Flow beads (GE Healthcare) for GST-tagged proteins at 4 °C following the manufacturer’s instructions. Eluted fractions were dialyzed into 150 mM NaCl, 50 mM Tris-HCl pH 7.5, 1 mM DTT and 5 mM MgCl_2_ and protein concentrations were determined by measuring the absorption at 280 nm. Nucleotide exchange to GTPγS or GDP was performed by the addition of 10-fold molar excess of GTPγS or GDP (Sigma Aldrich) and 5 mM EDTA to dialyzed proteins. After 30 min incubation at 37 °C, proteins were transferred on ice and exchange was quenched by the addition of 10 mM MgCl_2_ final concentration.

### Construction of a phage-displayed CRAF-RBD library

For phage library construction, Ras-binding domain (RBD) of human CRAF kinase (AA 55-131) was cloned into the phagemid pNE [21] and cysteines at position 81, 95 and 96 of the RBD were mutated to serines using site-directed mutagenesis [36]. Afterwards, two degenerate oligonucleotides were used to introduce mutations at 2 regions of the RBD gene by site-directed mutagenesis. Oligonucleotides were soft-randomized as described before [21, 22]. The nucleotide ratio was adjusted to 70 % of the wt nucleotide represented N1 = A, N2 = C, N3 = G and N4 = T and 10 % of each of the other three nucleotides.

Oligonucleotide 1: GAT GAC AAA AGC AAC N1N2N4 ATC N2N3N4 GTT N4N4N2 TTG CCG AAC N1N1N3 N2N1N1 N1N3N1 ACA N3N4N3 GTC N1N1N4 GTG CGA AAT GGA ATG

Oligonucleotide 2: CAT GAC TGC CTT ATG N1N1N1 N3N2N1 CTC N1N1N3 N3N4N3 N2N3N3 N3N3N2 CTG CAG CCA GAG TGC

The resulting library was used to electroporate *E. coli* SS320 cells using established methods resulting in 2.2×10^9^ independent RBD variants [37].

### RBD variant selection against activated HRAS

Library pool of phage displaying individual RBD variants were harvested by precipitation with PEG/NaCl (20 % PEG-8000 (w/v), 2.5 M NaCl) and resuspended in PBT buffer (1x PBS, 1 % BSA, 0.1 % Tween 20) supplemented with 5 mM MgCl_2_. Immobilization of GTPγS-loaded GST-tagged HRAS and subsequent binding selections were done essentially as described before [22], except that all buffers were supplemented with 5 mM MgCl_2_. In brief, 4 wells of a 96 well Maxisorp microtiter plate (NUNC) were coated with 100 µl of 2 µM GST-tagged GTPγS-loaded HRAS in dialysis buffer (150 mM NaCl, 50 mM Tris-HCl pH 7.5, 1 mM DTT and 5 mM MgCl_2_) over night at 4 °C. After blocking with PBT buffer for 1 h, phage library pool was added to each well and incubated for 1 hour at 4 °C. The plate was washed 8 times with cold PT buffer (1x PBS, 0.1 % Tween 20) and bound phage were eluted with 0.1 M HCl and immediately neutralized with 1.0 M Tris-HCl pH 8.0. Eluted phage were directly used to infect exponentially growing *E. coli* XL1-blue supplemented with helper phage M13K07 (NEB) and incubated over night at 37 °C. In each successive selection round, the selection stringency was increased by 2 additional washing steps and. To avoid unspecific binding towards the GST-tag, a GST counter-selection was performed starting with round 2. Additionally, after 3 rounds of selection, phage binding and washing was done at room temperature and the concentration of coated GST-tagged GTPγS-loaded HRAS was reduced to 0.5 µM. After five rounds of enrichment, individual RBD variants with improved binding properties towards active HRAS were identified by clonal phage ELISA as described [37]. In brief, phage displaying individual RBD variants were prepared from single colonies of bacteria harboring phagemids encoding RBDvs and transferred to 384-well Maxisorp plates immobilized with GST-tagged GTPγS-labelled HRAS (0.5 µM) and blocked with BSA, as described previously. As negative controls, wells were coated with GST or only blocked with BSA. After incubation, washing and developing, positive clones were further analysed by sequencing.

### Cloning of RBD variants

DNA fragments encoding the selected variants and the unmodified RBDwt were cloned into pDONR233 plasmids by Gateway cloning (Invitrogen) as described before [22]. Further recombination into the Gateway destination vectors pET53DEST, pDEST15 and pLDT-NT-HA was performed according to the manufacturer’s instructions (Invitrogen).

### RBDvs protein expression and purification

For competitive *in vitro* pull-down experiments and bio-layer interferometry measurements, His-tagged (from pET53DEST plasmids) and GST-tagged (from pGEX-6p-1 plasmids) RBD variants and RBDwt, Ras-association domain (RA) of RalGDS (AA 798-885) and GST alone were processed as Ras proteins (see above).

### Competitive *in vitro* pull-down experiment

The Ras/RBDwt competition assay was performed in 200 µl assay buffer (25 mM Tris-HCl pH 7.5, 150 mM NaCl, 1 % NP-40, 5 mM MgCl_2_ and 5 % glycerol) using 8 µg (1.15 µM) GST-tagged RBDwt or RalGDS-RA bound to GSH-sepharose beads. First, 5 µg (1.15 µM) of His-tagged KRAS loaded with GTPγS was incubated with His-tagged RBDwt or RBDvs with increasing concentrations (2.3 µg [1.15 µM], 5.8 µg [2.68 µM] and 23 µg [11.5 µM], which corresponds to a molar ratio of 1:1, 1:2.5 and 1:10 of GST-tagged RBDwt or RalGDS-RA:His-tagged RBDwt or RBDvs) for 30 min at 4 °C with end-over-end rotation. 8 µg of beads were added to the reaction mix and incubated for another 30 min at 4 °C. After 2 washes with assay buffer, beads were resuspended in 30 µl 2x SDS sample buffer and incubated for 5 min at 95 °C. Controls samples consisted of GTPγS- or GDP-loaded KRAS incubated with GST-RBDwt or GST-RalGDS-RA beads. Also, GTPγS-loaded KRAS was incubated with GST bound to beads. Samples were analyzed by immunoblot using anti-Ras antibody (#16117 ThermoFischer Scientific) and membrane was stained with Ponceau S as loading control.

### Bio-layer interferometry (BLI)

Kinetic binding assays were performed on Octet RED96 instrument (Pall ForteBio). Dialyzed proteins were supplemented with 0.1 % BSA and 0.02 % Tween 20. GST or GST-tagged RBDvs were immobilized onto anti-GST biosensors (Pall ForteBio) at a concentration of 2 µg/ml. Association was analyzed at concentrations starting from 1000 nM to 15.6 nM in 1:1 dilution steps of GTPγS- or GDP-loaded HRAS. Dissociation was measured in dialysis buffer (150 mM NaCl, 50 mM Tris-HCl pH 7.5, 1 mM DTT and 5 mM MgCl_2_) supplemented with 0.1 % BSA and 0.02 % Tween 20. Non-specific interaction of HRAS with biosensors was assayed using empty anti-GST sensors. Reference wells were subtracted from sample wells and a 1:1 global fitting model was used to determine *k*_on_, *k*_off_ and *K*_d_ values.

### Protein expression and purification for crystallography

RBDv1, RBDv12 and HRAS G12V (residues 1-166) were expressed as TEV protease cleavable GST-fusions using a modified pGEX-2T vector. Expression constructs were transformed into BL21-CodonPlus DE3-RIL bacteria (Agilent Technologies) for protein production. Bacterial expression was induced overnight at 18 °C with 0.5 mM IPTG and was performed in LB media. RBDvs bacterial pellets were resuspended in 50 mM HEPES pH 7.5, 300 mM NaCl, 5 % glycerol, 1 mM PMSF, 1 mM TCEP and lysed by homogenization. The lysate was cleared by centrifugation at 4 °C for 40 minutes at 18,000 g. Protein was bound to glutathione affinity resin (GE Healthcare), eluted by cleavage of the GST-tag with TEV, concentrated and then buffer exchanged by size exclusion chromatography (SEC) using a Superdex75 24 ml column (GE healthcare) equilibrated in 50 mM HEPES pH 7.5, 50 mM NaCl, 5 % glycerol, 10 mM MgCl_2_, 1 mM TCEP.

HRAS G12V bacterial pellets were resuspended in 50 mM HEPES pH7.5, 300 mM NaCl, 5 mM EDTA, 5 % glycerol, 1 mM PMSF, 1 mM TCEP and lysed by homogenization. The lysate was clarified by centrifugation at 4 °C for 40 minutes at 18,000 g. Protein was bound to glutathione affinity resin (GE Healthcare), washed with 50 mM HEPES pH 7.5, 300 mM NaCl, 25 mM imidazole, 5 % glycerol, 1 mM TCEP and eluted by cleavage of the GST-tag with TEV, concentrated and then buffer exchanged by SEC using a Superdex75 24 ml column equilibrated in 50 mM HEPES pH7.5, 300 mM NaCl, 5 mM EDTA, 5 % glycerol, 1 mM TCEP. GMP-PNP loading on HRAS G12V was carried out by incubating purified HRAS G12V with 10-fold excess of GMP-PNP for 80 min at 4 °C followed by the addition of 30-fold excess of MgCl_2_ for 120 min at 4 °C.

The RBDvs:HRAS G12V complexes were obtained by mixing RBDvs and GMP-PNP loaded HRAS G12V at equal molar ratio for 60 min at 4 °C followed by SEC using a Superdex75 24 ml column equilibrated in 50 mM HEPES pH 7.5, 50 mM NaCl, 10 mM MgCl_2_, 5 % glycerol, 1 mM TCEP. All fractions corresponding to co-elution of HRAS G12V with RBDvs were pooled and concentrated to 12.2 mg/ml, then flash frozen in liquid nitrogen. Protein concentration was estimated by UV-Vis absorption spectroscopy using a NanoDrop spectrophotometer (Thermo-Fisher Scientific).

### Protein crystallography, data collection and structural analysis

The RBDvs:HRAS G12V complexes were crystallized at 20 °C in sitting-drop by mixing 0.5 µL of complexes (450 μM, 12.2 mg/ml) with 0.5 µL of mother liquor of 0.1 M sodium cacodylate pH 6.5, 200 mM ammonium sulphate, 30 % PEG 4000 or mother liquor of 0.1 M sodium citrate pH 5.6, 200 mM ammonium sulphate, 30 % PEG 4000 for RBDv1:HRAS G12V and RBDv12:HRAS G12V, respectively. X-ray diffraction data were collected on a flash-frozen crystal cryo-protected in mother liquor containing 25 % glycerol at 100 K on station 24-ID-C, NE CAT beamline, Advanced Photon Source (APS). Data reduction was performed using XDS package [38]. The structure was solved by molecular replacement using PDB 4G0N [5] as a search model in Phaser [39]. Model building and refinement was performed using COOT [40], LORESTR [41] and REFMAC [42] from the CCP4 suite [43]. The data statistics and refinement details are reported in Supplementary Table 1.

### Cell culture

HCT 116 (#CCL-247) cells were purchased from ATCC and handled according to the supplier’s instructions. MIA PaCa-2, A549 and H1299 cell lines were cultured in DMEM (MIA PaCa-2 and A549) or RPMI-1640 (H1299) medium (both Gibco) supplemented with 10% (v/v) fetal bovine serum (Gibco). Cell lines were tested and found to be mycoplasma-free. Lenti viral particles were produced in HEK293T cells using established protocols [44]. Stable cell lines were constructed using lentiviral vectors (pLDT) encoding N-terminally HA-tagged RBDvs and wt under the control of a doxycycline inducible promotor by standard techniques [45]. The original pLDT plasmid was a gift from Jason Moffat (University of Toronto, The Donnelly Centre, 160 College St., Toronto). Two days after transduction, cell lines were selected with 2 µg/ml puromycin for HCT 116, MIA PaCa-2, A549 and H1299. For all experiments, 1 µg/ml doxycycline was used to induce expression of HA-tagged RBDvs or RBDwt.

### Immunoprecipitation for mass spectrometry

Two 10 cm dishes per construct (RBDwt, v1 and v12) were seeded with 2×10^6^/ml stable HCT 116 cells and RBDvs expression was induced 24 h later with doxycycline. After 24 h induction, cells were scraped into 1x PBS and washed twice with 1x PBS. After centrifugation, the cell pellets were resuspended in lysis buffer (25 mM Tris-HCl pH 7.5, 150 mM NaCl, 1 % NP-40, 5 mM MgCl_2_, 5 % glycerol and protease inhibitor cocktail [Roche]) and incubated for 20 min at 4 °C with end-over-end rotation. The lysates were centrifuged, supernatants were transferred to 20 µl pre-equilibrated anti-HA affinity matrix (Roche) and incubated for 40 min at 4 °C. Beads were washed 2 times with 500 µl lysis buffer. Elution was performed by the addition of 20 µl 2x SDS-sample buffer to the beads and incubation for 3 min at 95 °C. 5 µl of samples were analysed by immunoblotting using antibodies against endogenous Ras, HA-tag and GAPDH (#3965, #2999 and #8884, respectively; all from Cell Signaling Technology). Sample preparation for in-gel digest was done as described before [46]. In brief, supernatants were loaded onto SDS-polyacrylamide gel and the gel was stained using InstantBlue (Expedeon). Gel lanes were cut into pieces and subsequently washed, destained and dehydrated. Proteins were reduced with 10 mM dithiothreitol (DTT), alkylated with 55 mM iodoacetamide (IAA) and digested overnight with sequencing-grade Trypsin (Promega). Peptides were extracted using an increasing concentration of acetonitrile. Finally, peptides were concentrated and desalted using the Stop and Go Extraction (STAGE) technique [47].

### Liquid chromatography and mass spectrometry

A binary buffer system consisting of buffer A (0.1 % formic acid) and buffer B (80 % acetonitrile, 0.1 % formic acid) was used for peptide separation on an Easy-nLC 1200 (Thermo Fisher Scientific). This system was coupled *via* a nano electrospray ionization source to the quadrupole-based Q Exactive HF benchtop mass spectrometer [48]. Peptide elution from the in-house packed 18 cm (1.9 µm C18 Beads, Dr. Maisch, Germany) column was achieved by increasing the relative amount of B from 10 % to 38 % in a linear gradient within 23 min at a column temperature of 40 °C. Followed by an increase to 100 % B within 7 min, gradients were completed by a re-equilibration to 5 % B.

Q Exactive HF settings: MS spectra were acquired using 3E6 as an AGC target, a maximal injection time of 20 ms and a 60,000 resolution at 300 m/z. The mass spectrometer operated in a data dependent Top15 mode with subsequent acquisition of higher-energy collisional dissociation (HCD) fragmentation MS/MS spectra of the top 15 most intense peaks. Resolution for MS/MS spectra was set to 30,000 at 200 m/z, AGC target to 1E5, maximal injection time to 64 ms and the isolation window to 1.6 Th.

### Mass spectrometry data processing and analysis

All acquired raw files were processed using MaxQuant (1.5.3.30) [49] and the implemented Andromeda search engine [50]. For protein assignment, electrospray ionization-tandem mass spectrometry (ESI-MS/MS) fragmentation spectra were correlated with the Uniprot human database (v. 2016) including 4 manually added sequences of the RBD variants and KRAS G13D (RBDwt: MGYPYDVPDYAGQGPDPSTNSADITSLYK KAGFSNTIRVFLPNKQRTVVNVRNGMSLHDCLMKALKVRGLQPECCAVFRLLHEHKGKKARLDWNTDAASLIGEELQVDFL; RBDv1: MGYPYDVPDYAGQGPDPSTNSADIT SLYKKAGFSNTIRVLLPNQEWTVVKVRNGMSLHDSLMKALKRHGLQPESSAVFRLLHEHKGKKARLDWNTDAASLIGEELQVDFL; RBDv12: MGYPYDVPDYAGQGPDPST NSADITSLYKKAGFSNTIRVLLPNHERTVVKVRNGMSLHDSLMKALKRHGLQPESSAVFRLLHEHKGKKARLDWNTDAASLIGEELQVDFL; KRAS G13D: MTEYKLVVVGAGD VGKSALTIQLIQNHFVDEYDPTIEDSYRKQVVIDGETCLLDILDTAGQEEYSAMRDQYMRTGEGFLCVFAINNTKSFEDIHHYREQIKRVKDSEDVPMVLVGNKCDLPSRTVDTKQAQDLARSYGIPFIETSAKTRQGVDDAFYTLVREIRKHKEKMSKDGKKKKKKSKTKCVIM). Searches were performed with tryptic specifications and default settings for mass tolerances for MS and MS/MS spectra. Carbamidomethyl at cysteine residues was set as a fixed modification, while oxidation at methionine and acetylation at the N-terminus were defined as variable modifications. The minimal peptide length was set to seven amino acids and the false discovery rate for proteins and peptide-spectrum matches to 1 %. The match-between-run feature was used with a time window of 0.7 min. Relative label-free quantification of proteins was done using the MaxLFQ algorithm integrated into MaxQuant [51]. The minimum LFQ ratio count was set to 2 and the FastLFQ option was enabled.

For subsequent analysis, the Perseus software (1.5.3.0) [52] was used and first filtered for contaminants and reverse entries and proteins that were only identified by site. The LFQ intensities were logarithmized to base 2 and grouped into duplicates. To overcome the missing value problem in immunoprecipitation data, proteins that were quantified less than 2 times in one of the experimental groups were discarded from further analysis. Missing values were imputed column-wise by a down-shifted (median-1.8) Gaussian distribution mimicking the detection limit of the mass spectrometer.

### Metabolic activity assay

1000 cells of each stable cell line were seeded in a 96 well plate in duplicates. The following day, expression was induced by the addition of doxycycline. Corresponding control wells were not induced. Cell viability was assayed 120 h after induction using CellTiter-Glo (Promega). Luminescence signals of induced wells were normalized to uninduced well data.

### Immunoblotting

Stable cell lines were seeded in 6 well plates (1×10^5^ cells/well) and induced 24 h later with doxycycline (1µg/ml). Control cells were left uninduced. 24 h after induction, cells were washed twice with 1x PBS and lysed in 200 µl 2x SDS sample buffer. Whole cell lysates were analyzed by immunoblot using the indicated antibodies. For serum starvation experiments, medium was replaced upon induction with serum-free medium. 24 h after induction, cells were treated with 100 ng/ml EGF (Sigma) for 10 min before lysis.

### Antibodies

Antibodies used for western blot analysis were: pospho-ERK1/2 (p44/42 MAPK) (Thr202/Tyr204) (clone D13.14.4E #4370), ERK1/2 (p44/42 MAPK) (clone 137F5 #4695), phospho-AKT (Ser473) (clone D9E #4060), AKT (clone C67E7 #4691), GAPDH-HRP (clone D16H11 #8884), and HA-HRP (clone 6E2 #2999) were purchased from Cell Signaling Technologies.

### Analysis of cell growth by real-time cell analysis (RTCA)

Cell proliferation of stably transduced HCT 116 cells was assessed using xCELLigence real-time cell analysis (RTCA) SP (single plate) system (ACEA Biosciences). 1000 cells/well were seeded in a 96-well electronic microtiter plate (ACEA Biosciences) with 200 μl medium/well. After 24 h, cells were induced by doxycycline. Cell proliferation was monitored for 120 h. Experiments were performed as duplicates and repeated at least twice.

### Apoptosis analysis by flow cytometry

Stably transduced HCT 116 cells were seeded in 6-well plates (1×10^5^ cells/well) and induced 24 h later with doxycycline. Control cells were left uninduced. After 72 h, cells were washed once with 1x PBS. For analysis of apoptotic cells, Annexin V-FITC apoptosis detection kit (#ALX-850-020 Enzo) was used following the manufacturer’s protocol. Samples were analyzed on a FACSCanto II flow cytometer (BD Biosciences) and data were processed by FlowJo software (FlowJo, LLC). Gating was done based on viable and single cells that were identified on the basis of scatter morphology.

### Organoid cultures

Human tumor colon organoid samples were obtained from a published colorectal cancer organoid biobank [29]. Resection tissues were obtained with written informed consent, and following approval by the ethics committees of the Diakonessen Hospital, Utrecht. Tissue was obtained with written informed consent and following approval, according to the guidelines of the University Cancer Center (UCT), Frankfurt. Organoid cultures were established and maintained as described previously [53]. Tumor organoids were maintained in medium lacking Wnt3a. The organoid lines were transduced as described [54] with lentivirus expressing N-terminally HA-tagged RBDwt, RBDv1 or RBDv12 (as above). Three days after transduction, organoids were selected with 1 μg/mL puromycin in the culture medium. For cell viability assays, the cells were seeded in Matrigel either following mechanical dissociation or enzymatic single cell dispersal using TrypLE Express reagent (Gibco). 2 μg/mL doxycycline was added 1 day after seeding. After 3 days cell viability was measured using CellTiter-Glo reagent (Promega). All experiments were measured as technical triplicates and the experiments were repeated at least twice each.

For quantification of organoid size, light microscopy pictures were taken at a 2x resolution after 2-6 days of doxycycline exposure, as indicated. The diameter (µm) of all viable organoids in one picture of each condition was measured using ImageJ. Viable organoids were identified as refringent, while dark structures surrounded by cell debris were excluded from the analysis. The quantification was performed on two independent experiments.

### Statistical methods

Data are presented as the mean ± standard deviation (SD). The comparisons between RBDwt and RBDvs were made by an unpaired t test using GraphPad Prism software. For statistical analysis of the quantification results of organoid sizes, a Mann-Whitney U-test was performed using GraphPad Prism software. The level of significance was set at \**P*< 0.05, \*\**P*< 0.01 and \*\*\**P*< 0.005.

## Acknowledgments

This work was supported by LOEWE Ub-Net, Cluster of Excellence “Macromolecular Complexes” (DFG EXC115), and the Collaborative Research Center SFB 1177. We thank Mani Ravichandran from the Structural and Genomic Consortium (SGC) of Toronto for providing X-Ray crystallography screening kits. This work was supported by funds from an Impact Grant (704116 to F.S.) from the Canadian Cancer Society Research Institute and by operating funds (FDN143277 to F.S.) from the Canadian Institutes for Health Research. P.M. was supported by a TD Bank postdoctoral fellowship. F.S. holds a Canada Research Chair (Tier 1) in Structural Biology of Cell Signaling. K.R. is a Heisenberg professor of the DFG (RA1739/4-2). Diffraction work conducted at the Northeastern Collaborative Access Team beamlines was funded by the National Institute of General Medical Sciences from the National Institutes of Health (P41 GM103403) and by an NIH-ORIP HEI grant (S10 RR029205). We thank Nancy R. Gough (BioSerendipity, LLC) for constructive comments and editorial assistance. We thank Jason Moffat (University Toronto) for donation of inducible lentiviral plasmids.

## Author contributions

S.W. and A.E. designed the study, designed the RBD library, analyzed the data and wrote the manuscript. S.W. performed library construction and selection, *in vitro* competition experiments, binding experiments and all cellular experiments. P.M and I.K. performed structural studies and P.M., F.S., S.W. and A.E. analyzed structural data. S.W. performed Co-IP and MS and S.W., M.H. and A.E. analyzed MS data. K.R. contributed to the design, provided reagents and analyzed data. M.B.G. performed organoid experiments and M.B.G., H.F.F., S.W., H.C. and A.E. analyzed organoid data. M.K. analyzed data and provided reagents.

## Data and materials availability

All data is available in the main text or the supplementary materials. All data and material will be made available upon request.

