## Supplementary Figures and Tables for "Novel Conformation Specific Inhibitors of Activated GTPases reveal Ras-dependency of Patient-Derived Cancer Organoids"

**Titel page:**

### Supplementary Information

### Supplementary Figures 1 - 5

#### Supplementary Fig. 1

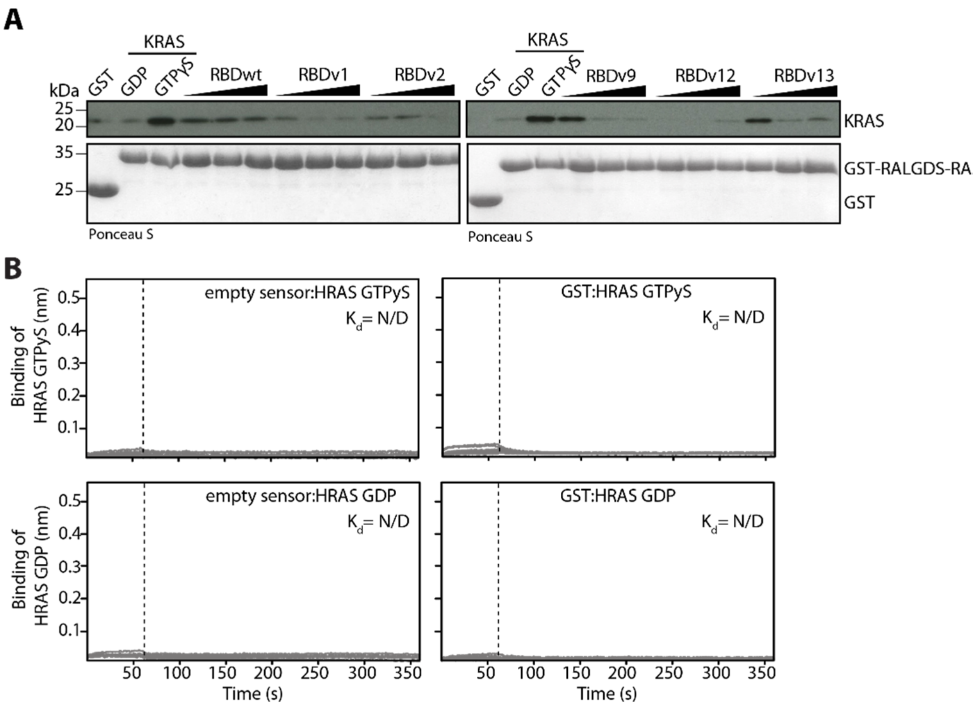

**C**

| RBD | $k_{on}$ [$10^5$x1/Ms] | $k_{off}$ [$10^{-3}$x1/s] | $K_d$ [nM] | Fold improvement |
|---|---|---|---|---|
| wt | 2.1 ± 0.2 | 12.8 ± 0.2 | 62.6 ± 4.8 | - |
| v1 | 3.8 ± 0.1 | 1.3 ± 0.1 | 3.4 ± 0.1 | **18** |
| v12 | 5.6 ± 0.2 | 1.4 ± 0.02 | 2.5 ± 0.1 | **25** |

**Supplementary Figure 1 - RBDvs outcompete RalGDS-Ras association domain (RA) binding to activated KRAS.**

(**A**) *In vitro* competition of increasing concentration of His-tagged RBDvs and RBDwt and GST-tagged RalGDS-RA immobilized on glutathione sepharose beads binding to His-tagged GTPγS-loaded KRAS. KRAS bound to beads was detected by immunoblot and the corresponding Ponceau S stained membrane is shown.

(**B**) Control experiments of GDP- or GTPγS-loaded HRAS and empty anti-GST biosensors measured by bio-layer interferometry. Concentrations of Ras ranged from 1 µM to 15.6 nM in a 1:1 dilution series. $K_d$ values for could not be calculated due to weak binding. N/D.=not determined.

(**C**) Binding constants for the BLI measurements from Fig. 1D. Values for on rate ($k_{on}$ [$10^5$x1/Ms]), off rate ($k_{off}$ [$10^{-3}$x1/s]), dissociation constant ($K_d$ [nM]) and fold improvement are shown.

### Supplementary Fig. 2

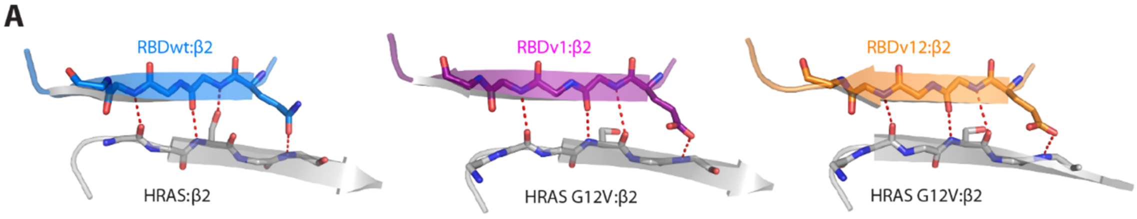

**Supplementary Figure 2 - RBDvs bind to HRAS through a canonical binding mode.** Detailed view of the canonical extended intermolecular β-sheet at the binding interface of the RBDwt or RBDvs with HRAS or HRAS G12V, respectively. Coloring and labeling are as in Fig. 2A.

### Supplementary Fig. 3

**A**

HA-Tag / attB1 / CRAF RBD (55-131)

```
RBDwt  MGYPYDVPDYAGQGPDPSTNSADITSLYKKAGFSNTIRVFLPNKQRTVVNVRNGMSLHDCLMKALKVRGLQPECCAVFRLLHEHKGKKARLDWNTDAASLIGEELQVDFL
RBDv1  MGYPYDVPDYAGQGPDPSTNSADITSLYKKAGFSNTIRVLLPNQEWTVVKVRNGMSLHDSLMKALKRHGLQPESSAVFRLLHEHKGKKARLDWNTDAASLIGEELQVDFL
RBDv12 MGYPYDVPDYAGQGPDPSTNSADITSLYKKAGFSNTIRVLLPNHERTVVKVRNGMSLHDSLMKALKRHGLQPESSAVFRLLHEHKGKKARLDWNTDAASLIGEELQVDFL
```

**B**

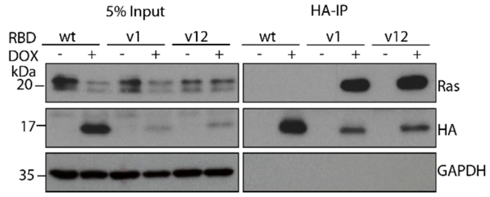

**Supplementary Figure 3 - Amino acid sequences of engineered RBDvs and RBDvs are binding to endogenous Ras in HCT 116 cells.**

(**A**) Amino acid sequences of HA-tagged RBDwt or RBDvs that were used in this study. Highlighted in bold (magenta and orange in RBDv1 and RBDv12, respectively) are those amino acids that differ from the RBDwt sequence. Underlined are the cysteine to serine mutations in RBDv1 and v12.

(**B**) Western blot of co-immunoprecipitation using anti-HA beads from lentiviral transduced HCT 116 cells stably expressing HA-tagged RBDwt, RBDv1 and RBDv12 upon induction with doxycycline (DOX) (1 µg/ml, 24h) using the indicated antibodies.

### Supplementary Fig. 4

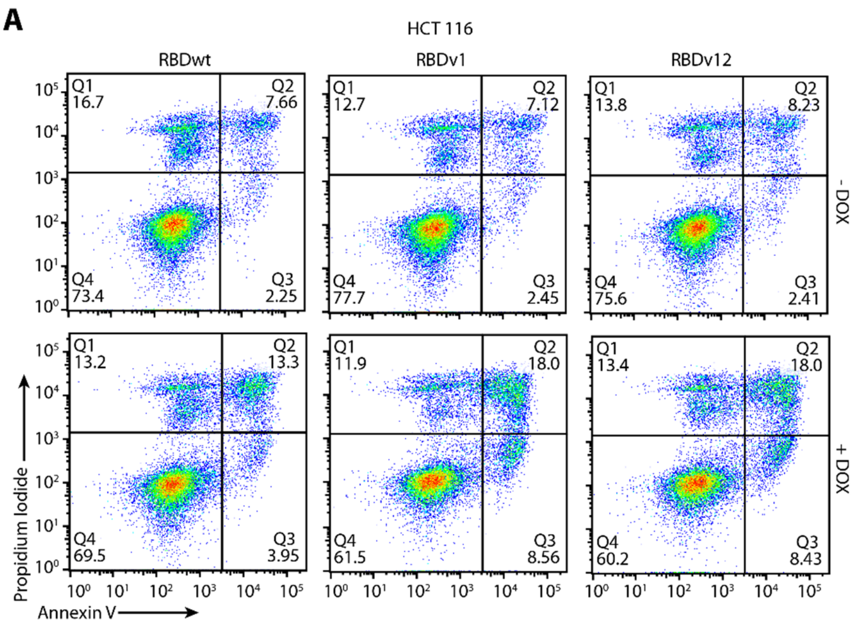

**Supplementary Figure 4 - RBDvs induce apoptosis in stable HCT 116 cells.** Representative flow cytometry analysis of HCT 116 cells stained with fluorophore labeled annexin V antibody and propidium iodide (PI) by flow cytometry in absence (-DOX) or presence (+DOX) of DOX (1 μg/ml, 72 h).

**Supplementary Fig. 5**

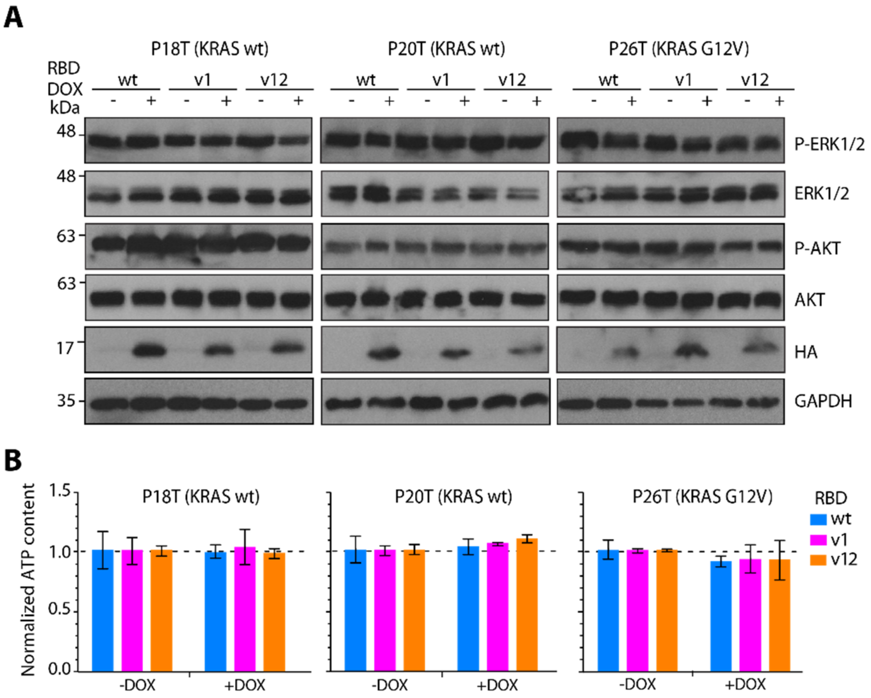

**Supplementary Figure 5 - Patient-derived colorectal cancer (CRC) organoids that were insensitive to RBDvs expression.**

(**A**) Immunoblot of whole cell lysates derived from indicated patient-derived CRC organoids stably transduced with lentivirus encoding HA-tagged RBDwt, RBDv1 and RBDv12 in absence (-) or presence (+) of DOX (2 µg/ml, 72 h). Cell lysates were analyzed using indicated antibodies.

(**B**) Cellular ATP content of organoid cultures used in (A) expressing RBDwt (blue), RBDv1 (magenta) and RBDv12 (orange) was measured in a luciferase mediated bioluminescence assay. Reduction of cellular ATP in presence of RBDvs (+DOX) was monitored after 72 h induction and normalized to the luminescence of non-induced control organoids (-DOX) (2 µg/ml, 72 h). Error bars correspond to ± SD of three technical replicates (n=3).

### Supplementary Tables 1 - 3

#### Supplementary Table 1: X-Ray data collection and refinement statistics.

| | **HRAS G12V:RBDv1** | **HRAS G12V:RBDv12** |
|---|---|---|
| **Data collection** | | |
| Space group | P 63 2 2 | P 63 2 2 |
| Cell dimensions | | |
| *a*, *b*, *c* (Å) | 91.62, 91.62, 151.41 | 91.76, 91.76, 151.59 |
| α, β, γ (°) | 90, 90, 120 | 90, 90 ,120 |
| Resolution (Å) | 151.4 - 2.9 (3.08 - 2.9) | 79.47 – 3.1 (3.31 – 3.1) |
| $R_{meas}$ | 0.14 (10.1) | 0.15 (5.31) |
| $CC_{1/2}$ | 99.9 (42.3) | 99.8 (43.9) |
| $I / \sigma I$ | 15.1 (0.4) | 11.0 (0.44) |
| Completeness (%) | 99.8 (98.7) | 97.9 (90.0) |
| Redundancy | 18.0 (18.7) | 12.3 (12.6) |
| **Refinement** | | |
| Resolution (Å) | 2.9 | 3.1 |
| $R_{work}$ / $R_{free}$ | 22.1 / 25.9 | 23.8 / 26.8 |
| No. atoms | | |
| Protein | 1779 | 1787 |
| Ligand/ion | 39 | 39 |
| Water | 8 | 1 |
| *B*-factors | | |
| Protein | 136.2 | 145.7 |
| Ligand/ion | 112.0 | 139.0 |
| Water | 100.0 | 109.3 |
| R.m.s. deviations | | |
| Bond lengths (Å) | 0.006 | 0.0086 |
| Bond angles (°) | 1.384 | 1.5068 |
| Ramachandran statistics | | |
| Residue in favoured regions (%) | 96 | 96.1 |
| Residue in allowed regions (%) | 4.0 | 3.5 |
| Residue in disallowed regions (%) | 0 | 0.4 |

**Supplementary Table 2: Dataset from co-immunoprecipitation and mass spectrometry analysis.**

See accompanying data set.

**Supplementary Table 3: Cell lines used in this study.**

| Name | Cell type | ATCC number | Ras mutation | Zygosity |
|---|---|---|---|---|
| HCT 116 | Human colon cancer cells | ATCC (#CCL-247) | KRAS G13D | Heterozygous |
| MIA PaCa-2 | Human pancreas cancer cells | ATCC (#CRL-1420) | KRAS G12C | Homozygous |
| A549 | Human lung cancer cells | ATCC (#CCL-185) | KRAS G12S | Homozygous |
| H1299 | Human lung cancer cells, derived from metastatic lymph node | ATCC (#CRL-5803) | NRAS Q61K | Heterozygous |
